# Statistics of spatial rotations and projection directions considering molecular symmetry in 3D electron cryo-microscopy

**DOI:** 10.1101/2020.08.24.264952

**Authors:** Qi Zhang, Hai Lin, Mingxu Hu

**Author notes:** Correspondence should be addressed to M.H. and H.L.

## Abstract

Electron cryo-microscopy (cryoEM) three-dimensional (3D) reconstruction is based on estimations of orientations of projection images or 3D volumes. It is common that the macromolecules studied by cryoEM have molecular symmetry, which, unfortunately, has not been taken into consideration by any statistics for either spatial rotations or projection directions at this point. Meanwhile, there are growing needs to adopt advanced statistical methods, and further, modern machine learning techniques in cryoEM. Since those methodologies are built heavily upon statistical learning cornerstones, the absence of their domain-specific statistical justification limits their applications in cryoEM. In this research, based on the concept of non-unique-games (NUG), we propose two key statistical measurements, the mean and the variance, of both spatial rotations and projection directions when molecular symmetry is considered. Such methods are implemented in the open-source python package pySymStat.

## 1 Introduction

In the three-dimensional (3D) reconstruction of electron cryo-microscopy (cryoEM), particle images are projections of biological samples at random orientations. Therefore, the analysis of such orientations is the key step in the reconstruction of 3D density maps[1, 2, 3, 4, 5, 6, 7, 8]. Accordingly, the analysis of both spatial rotations and projection directions is fundamentally important for cryoEM. In the cryoEM field, it is common that the macromolecules studied have molecular symmetry[9, 10]. However, the true topology induced by the molecular symmetry is currently neglected as there are no statistics considering molecular symmetry.

Meanwhile, there are growing demands to adopt advanced statistical methods, and further, machine learning techniques[11, 12] in cryoEM. Modern machine learning practices, including deep learning, are built upon statistical learning cornerstones. Therefore, such absence of statistics tools limits the applications of advanced statistical methods and machine learning techniques in cryoEM, in the circumstances that the macromolecules studied have molecular symmetry. Those methodologies would likely fail if true topology induced by the molecular symmetry is neglected, especially when the order of molecular symmetry is high.

In this study, we propose statistics of both spatial rotations and projection directions which take molecular symmetry into consideration. Such methods are realized in the open-source python package pySymStat. Our solution starts with representing spatial rotations using unit quaternions. We then estimate the mean and variance in a non-unique-games (NUG) framework, which is devised for solving a certain type of optimization problems defined over a locally compact group. The solution to our optimization problem is the convex combination[13] of all possible estimates, therefore we utilize a leaderboard algorithm to recover the final true result, i.e., the global optimal solution, from the NUG estimates.

The unit quaternion[14, 15, 16] is widely used for describing rotations. Section 2 gives a brief overview of this concept as well as clarifies the notations used in this work. In Section 3, we define distance, variance, and mean in the context of considering molecular symmetry. In Section 4, we demonstrate how such variance and mean problems can be solved by a method developed based on NUG[17, 18]. In Section 5, the performance of the proposed methods is analyzed. After determining the distance, variance, and mean, clustering is introduced in Section 6. The potential applications of the proposed methods are discussed in Section 7.

## 2 Unit quaternion description

The unit quaternion description of rotation has been well studied and applied in many fields, including robotics[19, 20, 21, 22], aviation[23, 24, 25, 26], computational vision[27, 28, 29], as well as cryoEM[30, 31, 32, 33, 34, 7, 35, 5, 36, 37, 38, 39, 40, 41, 42, 43, 44, 45]. In the present work, we give a very brief introduction of the unit quaternion description of spatial rotations. For general introductions of quaternion, we refer the reader to [15, 16, 46, 47].

Notations in this article are specified according to the following rules. The unit quaternion in vector form, which is a unit four-dimensional (4D) vector, is denoted by a bold letter, e.g., **q**. The molecular symmetry group is denoted by *G*. An element in the molecular symmetry group *G* is denoted by **g**. Such an element is also in the unit quaternion description. Meanwhile, the number of elements in *G* is denoted by |*G*|. The set containing all 3D spatial rotations is denoted by *SO*(3).

A quaternion is an extension of a complex number, composed of a real part and three imaginary parts. The quaternion is commonly viewed as a 4D vector. If its norm equals 1, i.e., |**q**| = 1, the quaternion is called a unit quaternion. Quaternion multiplication follows special rules, whose definition can be found in [15, 16]. The quaternion multiplication operator is denoted by ⊗ in the present work.

The rotation of a 3D vector 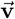 to another vector 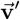 can be achieved by quaternion multiplication. By viewing 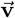 as 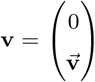 in quaternion form, the rotation induced by a unit quaternion is defined by[15]

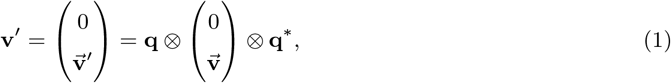

where **q**\* is the conjugate[47, 15, 16] of **q**. Therefore, **q** is a description of such spatial rotation. Such rotation is also denoted by

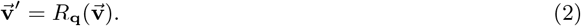

For sequential rotations, such as a spatial rotation **q**_1_ followed by a spatial rotation **q**_2_, the overall rotation is calculated as **q**_2_ ⊗ **q**_1_[15]. When molecular symmetry is not considered, the distance between the two spatial rotations **q**_1_ and **q**_2_ can be defined by[47].

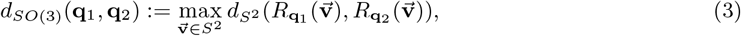

where 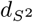 is the natural distance in the unit sphere *S*^2^.

Meanwhile, there are several different definitions of the projection direction in the cryoEM field. Some of them, however, are ambiguous. Hence, we explicitly define the projection direction as follows, in the present work. We denote the operator of projection for a spatial rotation **q** as 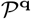. In other words, 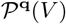 stands for the image obtained as follows. The map *V* is rotated by spatial rotation **q**; then, the rotated map is projected along the *Z* axis. The defining of projection direction 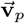 can be achieved by defining the set of projected images along 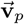, which is denoted by 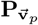. This set is defined by

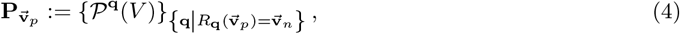

where 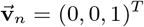 is the unit vector along the *Z* axis. In other words, every projection image in this set is projected along the *Z* axis from a rotated volume. Such a rotation transforms 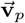 to 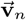. For two projection directions, 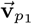 and 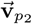, they are equal if and only if their corresponding sets of projected images are equal, i.e., 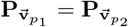. The definition of the distance between the two projection directions 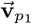 and 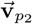 is assigned by the natural distance in the unit sphere *S*^2^, when molecular symmetry is not taken into consideration.

## 3 Definitions of distance, variance, and mean considering molecular symmetry

In this section, we redefine two most basic statistical measurements, the mean and the variance, in the context of considering molecular symmetry. We first define the equivalence relations on the sets of spatial rotations and projection directions, respectively (Section 3.1). Their interset distances are then used to measure the distances between two spatial rotations or two spatial rotations, respectively (Section 3.2), based on which we define the mean and the variance accordingly (Section 3.3).

### 3.1 Equivalence relations considering molecular symmetry

The symmetry property of a molecule can be represented by a molecular symmetry group. For a molecule with symmetry *G*, the spatial rotation **q** is equivalent to **q** ⊗ **g**_*m*_ for any **g**_*m*_ in a molecular symmetry group *G*. This is proven in Appendix I. We denote all of these equivalent spatial rotations of **q** as **q** ⊗ *G*, defined by

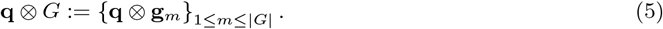

In mathematical terms, **q** ⊗ *G* is a left coset[48, 49] of *G* in *SO*(3). An element in the set of equivalent spatial rotations **q** ⊗ *G*, i.e., **q** ⊗ **g**_*m*_, is referred to as a representative[48, 49] of this set.

Meanwhile, when the molecular symmetry *G* is taken into consideration, the projection direction 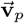 is equivalent to 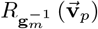 for any **g**_*m*_ in *G*; this is proven in Appendix II. Here, an inverse operation of **g**_*m*_ exists for the completeness of the concept. 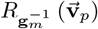 is the equivalent projection direction generated by rotating the map *V* by the symmetry element **g**_*m*_ (see Appendix II).

We denote all of these equivalent projection directions of 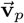 as 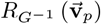, defined by

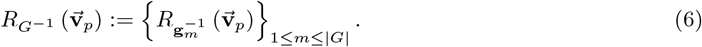

An element in the set of equivalent projection direction 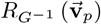, i.e., 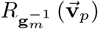, is referred to as a representative of this set.

For a set of spatial rotations {**q**_*i*_}_1≤*i*≤*N*_ and a set of projection directions 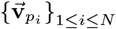, their statistical analyses are actually performed on {**q**_*i*_ ⊗ *G*}_1≤*i*≤*N*_ and 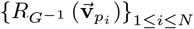 respectively, when the molecular symmetry *G* is taken into consideration.

### 3.2 Distance considering molecular symmetry

Definitions of the distance between two sets of equivalent spatial rotations and the distance between two sets of equivalent projection directions are the prerequisites of considering the statistics of spatial rotations and projection directions, respectively, when molecular symmetry is taken into consideration.

It is a common approach to define the distance between two sets by the distance between their representatives[50]. For spatial rotations, we define the distance between **q**_1_ ⊗ *G* and **q**_2_ ⊗ *G* by

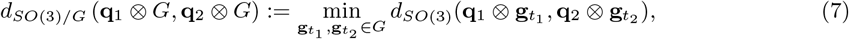

which is the shortest distance in *SO*(3) by selecting representatives 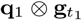 and 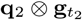 from **q**_1_ ⊗ *G* and **q**_2_ ⊗ *G*, respectively.

Similar to the distance definition of two spatial rotations, we define the distance between two sets of equivalent projection directions 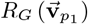 and 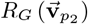 by

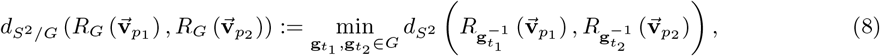

which is the shortest distance in *S*^2^ by selecting representatives 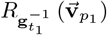 and 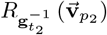 from 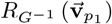 and 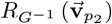, respectively. 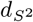 is the natural distance in the unit sphere *S*^2^.

These definitions, i.e., Equation (7) and Equation (8), both satisfy all of the requirements of the distance definition in mathematics (see Appendix III and Appendix IV, respectively), namely, the symmetry property, the positive definiteness property, and the triangle inequality property.

### 3.3 Mean and variance considering molecular symmetry

In the same spirit of the distance definition, we define the variance of spatial projections {**q**_*i*_ ⊗ *G*}_1≤*i*≤*N*_ and projection directions 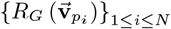, respectively as

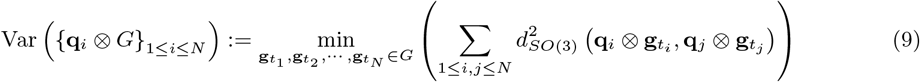

and

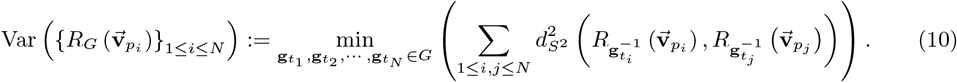

In other words, for spatial rotations, a representative 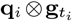 is chosen from each set of equivalent spatial rotations **q**_*i*_ ⊗ *G*, such that all representatives are as close as possible overall. The projection direction follows the same principle. Furthermore, we define the mean of spatial rotations {**q**_*i*_ ⊗ *G*}_1≤*i*≤*N*_ and projection directions 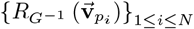, respectively as

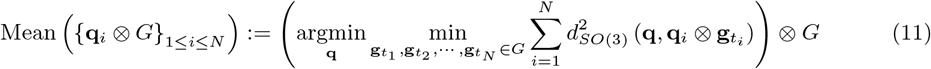

and

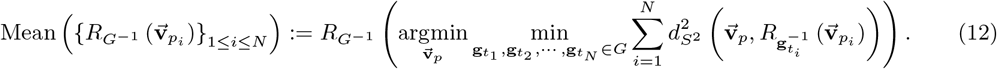

In other words, for spatial rotations, the mean of {**q**_*i*_ ⊗ *G*}_1≤*i*≤*N*_ is defined by the spatial rotation closest to {**q**_*i*_ ⊗ *G*}_1≤*i*≤*N*_. The mean of the projection direction follows the same principle.

The problems of determining the variance and the mean are in some sense equivalent. For spatial rotations, the mean definition by Equation (11) can be approximated by (see Appendix V)

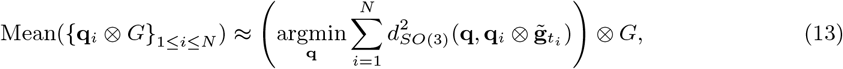

where

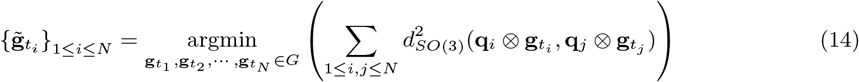

and 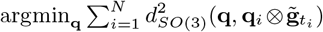 is solvable by an established method[30] once 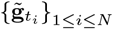 are determined. Meanwhile, for projection directions, the mean definition by Equation (12) can be approximated by (see Appendix VI)

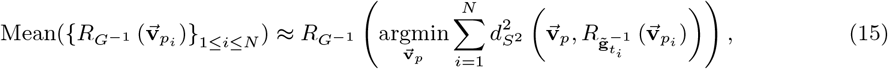

where

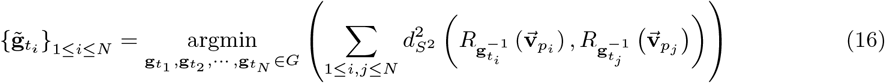

and 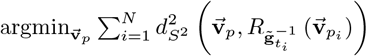 is also solvable by an established method[51] once 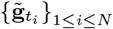 are determined.

In other words, the representatives chosen in the determination of the variances, i.e., Equation (9) and (10), determine 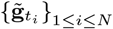. The means of spatial rotations and projection directions can be approximated by Equation (13) and Equation (15), by using 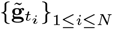.

In conclusion, choosing the overall-closest representatives, i.e., 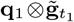,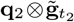, …, 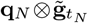 for spatial rotations, and 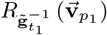, 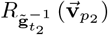, … , 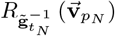 for projection directions, where 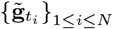 are selected by Equation (14) and Equation (16), is the key step in computing the statistics of spatial rotations and projection directions, respectively, when molecular symmetry is taken into consideration. In other words, once Equation (14) and Equation (16) are solved, the variance and the mean of {**q**_*i*_ ⊗ *G*}_1≤*i*≤*N*_ and 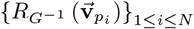 can be calculated, respectively.

## 4 NUG and the determination of representatives

In this section, we present an approximation algorithm for determining 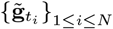 in both Equation (14) and Equation (16), which is equivalent to the variance and the mean determination, as discussed above. This approximation algorithm is based on the framework of non-unique games (NUG), developed by Lederman and Singer[17, 18]. Therefore, we refer to this algorithm as the NUG approach. The 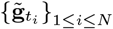 determination in both Equation (14) and Equation (16) is first translated into a NUG problem (see Section 4.1). Then, by making use of representation theory[52, 53, 54, 55, 56, 57, 58, 59, 60, 61, 62, 17, 18, 63] (see Section 4.2 for a brief overview), the NUG problem is relaxed into a semi-positive definite programming (SDP) problem[64, 65] (see Section 4.3 and Section 4.4), which can be solved by a convex optimization technique[64, 13]. However, the solution of the SDP problem is a “relaxation” of the NUG problem. In other words, the constraint in the SDP problem is looser than that of the NUG problem. Therefore, in the present work, we propose a leaderboard algorithm for “re-constraining” the SDP solution into a proper solution for the NUG problem (see Section 4.5). Currently, such algorithms are mainly adopted as leaderboard peptide sequencing in bioinformatics[66, 67].

### 4.1 Determination of representatives is a NUG problem

The original NUG problem is defined by the following optimization problem

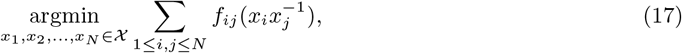

where 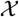 is a given locally compact group, and {*f*_*ij*_}_1≤*i,j*≤*N*_ are *N*^2^ given real valued functions on 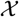.

The 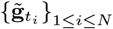 determination for both spatial rotation statistics, i.e., Equation (14), and projection direction statistics, i.e., Equation (16), can be derived in the same form (see Appendix VII and Appendix VIII)

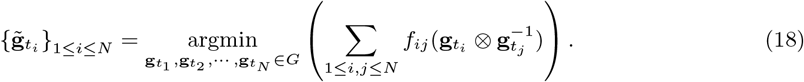

In the above equation, 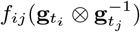 is assigned as (see Appendix VII),

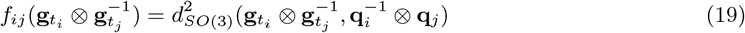

for the representative determination of spatial rotations. Meanwhile, for the representative determination of projection directions, 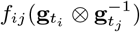 is assigned as (see Appendix VIII),

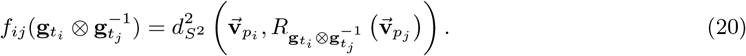

Moreover, the molecular symmetry group *G* is a finite group. Hence, the determination of representatives meets all of the requirements of being a NUG problem. In the following, we solve this NUG problem for determining the variance and the mean for both spatial rotations and projection directions.

### 4.2 Recall of representation theory

As in the NUG problem, the variables in the optimization are elements of the molecular symmetry group *G*. However, it is often difficult to optimize over a finite group. Proposed by Singer and Lederman[17, 18], such problems can be converted into optimizations over matrices using representation theory[52, 53, 54, 55, 56, 57, 58, 59, 60, 61, 62, 63]. Representation theory provides a general Fourier transform between a group and matrices, making such conversions possible.

We would like to briefly present some useful concepts and methods of representation theory in this section. As the molecular symmetry group *G* is considered in the present work and it is a finite group, this overview is limited to representation theory of the finite group[59, 60, 61, 62, 63].

For a finite group *G*, it corresponds to several intrinsically different irreducible representations. The number of irreducible representations of *G* is denoted by *K*. The irreducible representations of *G* are denoted by {***ρ***_*k*_}_1≤*k*≤*K*_. The irreducible representation ***ρ***_*k*_ is a function, which maps every element in a group *G* to a real or complex *d*_*k*_ × *d*_*k*_ unitary matrix, where *d*_*k*_ is the dimensionality of the *k*-th irreducible representation. When *G* is given, its irreducible representations, i.e., {***ρ***_*k*_(**g**_*m*_)}_1≤*k*≤*K*_ are known. The irreducible representations have the property described below.

By the Peter-Weyl theorem[68], a function *f* : *G* → ℝ has a general Fourier transformation, as follows. The *k*-th Fourier coefficient is defined by

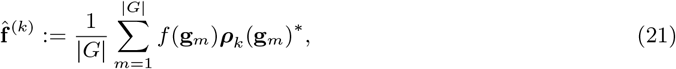

where ***ρ***_*k*_(**g**_*m*_)* is the conjugate transpose of ***ρ***_*k*_(**g**_*m*_). Then, the general Fourier expansion of *f* is

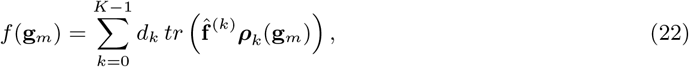

where 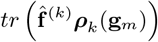 is the trace of the matrix 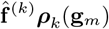.

### 4.3 Matrix form of a NUG by representation theory

By a general Fourier transformation using representation theory, introduced as Equation (21) and Equation (22), the NUG problem, i.e., Equation (14), is converted into the matrix form as (see Appendix IX)

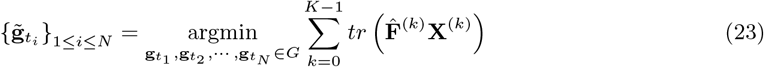

where

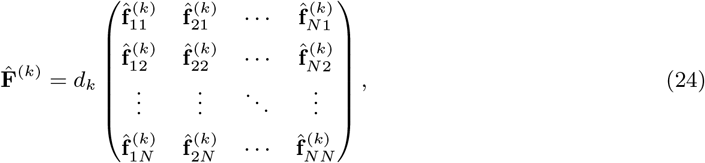

in which

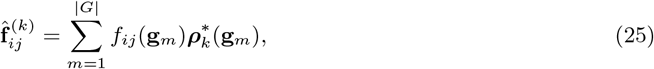

and

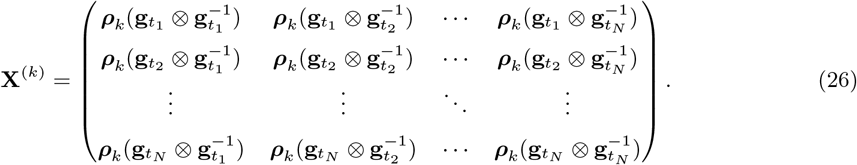

### 4.4 Convex relaxation of NUG to SDP

In the matrix form of a NUG problem, i.e., Equation (23), 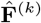 is known. As shown by Equation (25), the element of 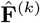, i.e., 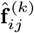, is determined by the given symmetry group *G* and the values of function *f*_*ij*_. In both cases (spatial rotations and projection directions), the values of the function *f_ij_* are pre-determined by {**q**_*i*_}_1≤*i*≤*N*_ and 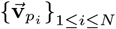 by Equation (19) and Equation (20), respectively. Meanwhile, **X**^(*k*)^ is composed of the variables, i.e., 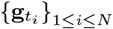, in the optimization problem. Thus, Equation (23) is an optimization problem over the variables {**X**^(*k*)^}_0≤*k*≤*K*−1_, subject to Equation (26) as constraints.

However, these constraints make the defined domain in this optimization problem in general non-convex. Thus, convex optimization techniques are not suitable, leading to the difficulty of determining a global optimal solution. However, such matrices {**X**^(*k*)^}_1≤*k*≤*K*_ can be relaxed into their convex hull[18, 69, 70] (see Appendix X), converting this NUG problem to an SDP problem. It is a classic problem to determine the solution of an SDP problem. In convex optimization, the global optimal solution is guaranteed. One can obtain the solution using open-source packages[71, 72, 73, 74]. Thus, instead of solving 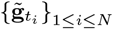, the solution matrices 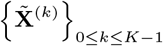 are obtained. Such solution matrices 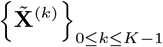 are in the convex hull of the matrices defined in Equation (26).

### 4.5 Recovery of representatives from the solution of SDP

As the constraint in the SDP problem is looser than that in the NUG problem, the solution matrices of the SDP problem may no longer comply with the constraints of the NUG problem, i.e., Equation (26). In other words, 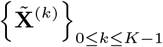, i.e., the solution matrices of the SDP problem, may violate Equation (26). However, determination of representatives, i.e, determination of 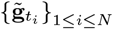, is needed for calculating mean and variance of rotations considering molecular symmetry. Thus, a recovery algorithm is needed for “re-constraining” 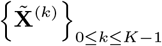 to obtain 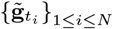.

We propose this recovery algorithm based on the following. Each block of the matrices {**X**^(*k*)^}_0≤*k*≤*K*−1_ i.e., 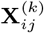, is constrained as

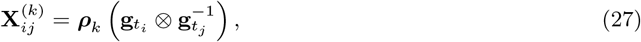

shown in Equation (26). Meanwhile, such a NUG problem is relaxed into an SDP problem by relaxing the above constraints into their convex hull. Based on the nature of a convex hull, the solution matrices of an SDP problem are the convex combinations of its constraints. In other words, each block 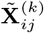 of 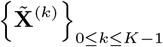 is the convex combination of ***ρ***_*k*_(**g**_*m*_) as

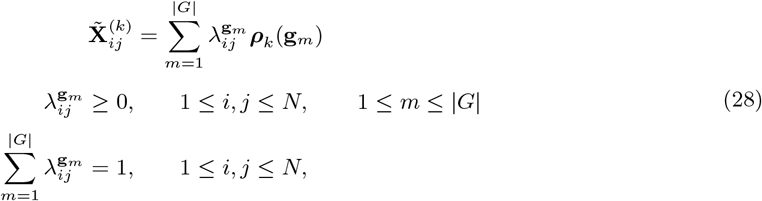

where 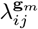 is the coefficient of the convex combination. 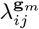 is determined as

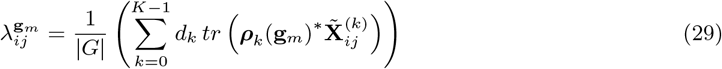

by the Peter-Weyl theorem[68], i.e., Equation (21) and Equation (22).

Comparing the constraints of the block matrices of the NUG problem, i.e., Equation (27), and the solution block matrices of the SDP problem as the convex combinations of ***ρ***_*k*_(**g**_*m*_), i.e., Equation (28), the convex combination coefficient 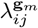 can be considered as the “probability” of 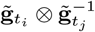 being **g**_*m*_.

For determining 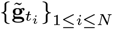 in the original NUG problem, *N* − 1 pairs of relations between two of them are required. In other words, once proper *N* −1 relations, i.e., 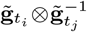, are determined, 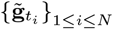 can also be determined. Meanwhile, as previously stated, 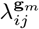 indicates the “probability” of 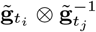 being **g**_*m*_. There are in total *N* × *N* × |*G*| guesses of every 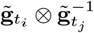 being every **g**_*m*_ in *G*; each has the “probability” 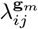. Hence, these guesses are redundant for solving 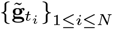.

In the present work, we propose a leaderboard algorithm for recovering 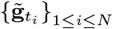 from such redundant guesses. This algorithm follows the following three principles.

The first principle of this algorithm is sequentially determining each 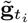. As *G* is a group, the following property holds. For all 1 ≤ *t*_*i*_, *t*_*j*_, *t*_*k*_ ≤ *N*, the relations between 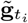 and 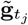, i.e., 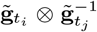, and the relation between 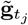 and 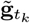, i.e., 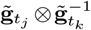, are recovered. In addition, the relation between 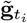 and 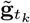, i.e., 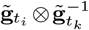, is also recovered. Therefore, this sequential determination of 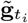 is a classic union-find[75]. In this union-find, if the relation between 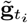 and 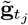 is decided, 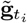 is connected to 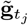. Once every 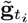 is connected to any other 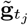, then 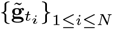 are determined.

The second principle of this algorithm is as follows. During sequentially deciding 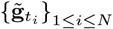, the relations between 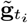 and 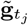 are sequentially determined in the descending order of their certainty. For example, 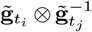 and 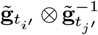 are two relations. As previously stated, the “probability” of 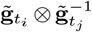 being **g**_*m*_ is 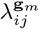. If the greatest value in 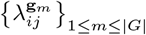 is greater than that in 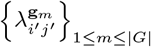, then the relation between 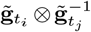 is more certain than that between 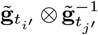. Therefore, the relation between 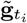 and 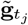 must be determined before the relation between 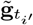 and 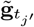. The reason of adopting this principle is as follows. As previously stated, it is redundant for solving 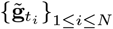 by *N* × *N* × |*G*| guesses of all 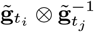. By deciding relations with high certainty first, high quality information is used for solving the problem which contributes to the high confidence of the recovery.

The third principle of this algorithm is as follows. For the relation 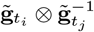, instead of determining it solely by **g**_*m*_ of the single greatest value of 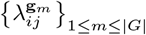, several trials are examined. These trials are based on multiple greatest values of 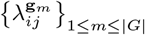. During the sequential determination of each 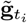, multiple sets of trials are tracked; these are selected by partial cost (see Appendix XI). Ultimately, the most qualified set of trials is selected as the final solution. This third principle makes the proposed algorithm in this work tally with the leaderboard algorithm.

## 5 Performance of the NUG approach

In this section, we empirically demonstrate that the NUG approach is capable of reaching the global optimal solution. The capability of solving Equation (14), i.e., selection of representatives for solving the variance and the mean of spatial rotations, is presented below (see Section 5.1). Moreover, since currently there are no other approximation algorithms for solving this problem, we have benchmarked our proposed NUG approach against a few intuitive alternatives, and have shown that the proposed NUG outperforms them all(see Section 5.2). For solving variance and mean of projection directions, the performance is similar, as Equation (16) has a similar construction to Equation (14).

### 5.1 Capability of reaching global optimal solution

Currently, there is no algorithm for obtaining a global optimal solution, other than the brute force approach. In the brute force approach, for each 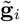, there are |*G*| potential **g**_*m*_ to be considered. Therefore, there are |*G*|^*N*^ trials to be examined. In other words, the computation cost grows exponentially as the number of spatial rotations grows. Hence, the global optimal is only practically solvable by the brute force approach when *N* is limited.

Several molecular symmetries are tested. For 3-fold cyclic (*C*_3_), 7-fold dihedral (*D*_7_), and tetrahedral (*T*) symmetries, *N* = 5 is used for performance tests due to the computational limitations of a modern workstation. Meanwhile, for octahedral (*O*) and icosahedral (*I*) symmetries, *N* is further limited to 3.

Such tests demonstrate that, in almost all cases, the NUG approach reaches a global optimal solution (see Table 1). Moreover, the capability of reaching a global optimal solution increases as the spatial rotations concentrate. Spatial rotations with four angular precision values[47] are tested. For the angular precision values higher than 7.9°, all cases tested reach a global optimal (see Table 1). Moreover, for cases in which global optimal solutions are not reached, the cost gaps between the obtained solution and the global optimal are small. For all cases tested, the maximum gap is approximately 5% (see Table 1).

**Table 1:** Performance of the NUG approach, compared with a intuitive approach

| Method | Symmetry | $\epsilon = 90^\circ^*$ | | $\epsilon \approx 36^\circ$ | | $\epsilon \approx 7.9^\circ$ | | $\epsilon \approx 1.6^\circ$ | |
| --- | --- | --- | --- | --- | --- | --- | --- | --- | --- |
| | | $nW/nT^{**}$ | $\delta^{***}$ | $nW/nT$ | $\delta$ | $nW/nT$ | $\delta$ | $nW/nT$ | $\delta$ |
| NUG approach | $C_3$ | 1/1000 | 1.78% | 1/1000 | 1.15% | 0/1000 | 0 | 0/1000 | 0 |
| | $D_7$ | 3/1000 | 5.07% | 1/1000 | 0.005% | 0/1000 | 0 | 0/1000 | 0 |
| | $T$ | 0/1000 | 0 | 0/1000 | 0 | 0/1000 | 0 | 0/1000 | 0 |
| | $O$ | 1/1000 | 0.002% | 0/1000 | 0 | 0/1000 | 0 | 0/1000 | 0 |
| | $I$ | 0/1000 | 0 | 1/1000 | 1.74% | 0/1000 | 0 | 0/1000 | 0 |
| intuitive approach | $C_3$ | 533/1000 | 23.0% | 260/1000 | 33.7% | 27/1000 | 8.6% | 1/1000 | 18.6% |
| | $D_7$ | 511/1000 | 33.7% | 441/1000 | 39.2% | 90/1000 | 44.3% | 5/1000 | 10.6% |
| | $T$ | 444/1000 | 44.8% | 367/1000 | 51.2% | 57/1000 | 43.9% | 2/1000 | 3.27% |
| | $O$ | 485/1000 | 59.0% | 429/1000 | 55.2% | 78/1000 | 49.6% | 6/1000 | 36.6% |
| | $I$ | 177/1000 | 70.0% | 177/1000 | 75.6% | 47/1000 | 60.5% | 3/1000 | 30.1% |

\* $\epsilon$ stands for the angular precision[47] of the testing spatial rotations.
\*\* $nW$ stands for the number of cases that fail to reach a global optimum. $nT$ stands for the number of cases tested.
\*\*\* $\delta$ stands for the maximum relative cost gap between the obtained solutions and the global optimal.

### 5.2 Comparison with an intuitive approach

The NUG approach is compared with an intuitive approach we design, as currently there is no other approximation algorithm available for solving Equation (14). We design several intuitive approaches, and one intuitive approach performs better than the others. The superior intuitive approach is described in Appendix XII.

However, compared to the NUG approach, the intuitive approach performs rather poorly (see Table 1). For spatial rotations in a uniform distribution, i.e., angular precision equals to 90°, 177 ~ 533 out of 1000 tested cases do not reach a global optimal, under conditions of various molecular symmetries. Moreover, the cost gaps between the obtained solution and the global optimal solution varies from 23.0% to 70.0% with molecular symmetries. Such incapability to reach a global optimal is alleviated as the spatial rotations concentrate. However, even when angular precision is 1.6°, there are cases which fail to reach global optimal solutions. Especially, the maximum cost gaps between the obtained solutions and the global optimal remain large even though the spatial rotations concentrate.

## 6 Clustering considering molecular symmetry

As the distance is defined and the mean can be computed, the clustering of both spatial rotations and projection directions can be achieved by several algorithms, e.g., classic k-means clustering[76]. Due to the similar construction of the distance definition and the mean calculation between the spatial rotations and the project directions, we can restrict our exposition to projection directions; equivalent results hold for spatial rotations.

As an example, we classified projection directions into five classes with several molecular symmetries (Figure 1). One may notice that, for some classes, the regions seem to be not connected. For example, consider the classes indicated by blue and turquoise in *C*_3_, and the classes indicated by turquoise and red in *D*_7_. However, when molecular symmetry is considered, these regions are in fact connected. Taking the red class with *D*_7_ symmetry as an example, although the right small region seems apart from the left large region, it can be rotated by a symmetry operation and then be attached to the left large region.

**Figure 1:**
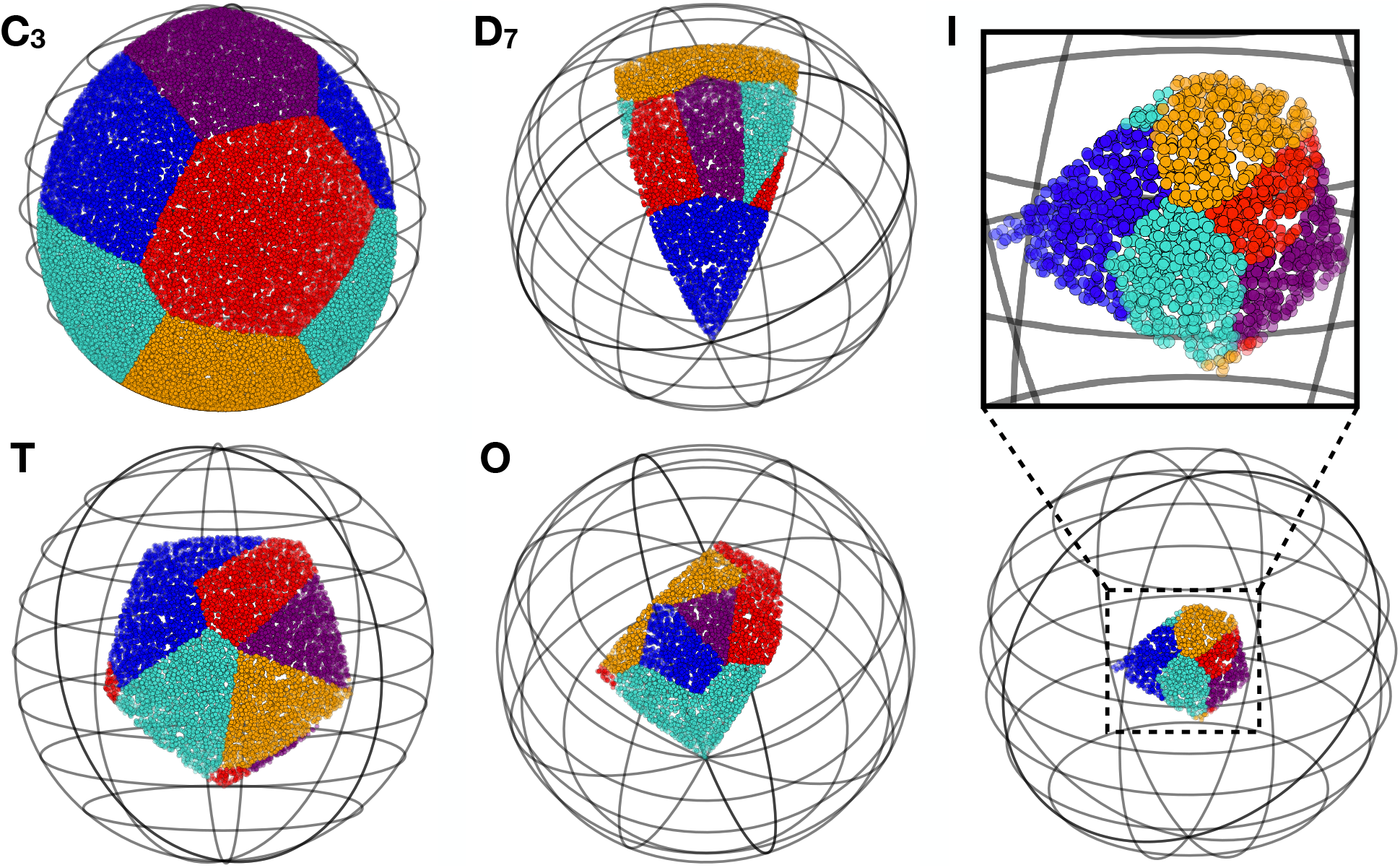
Clustering of projection directions considering molecular symmetries. With several molecular symmetries, the projection directions are classified into five classes; these are distinguished by color. For presentation purposes, in each set of equivalent projection directions 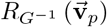, an element within a given fundamental domain is selected for display. The spheres are rotated in order to place projection directions towards the reader.

In contrast, the traditional method of projection directions clustering with molecular symmetry is as follows. A fundamental domain of *S*^2^*/G* is selected, and the projection directions are rotated by a symmetry operation into this domain. Clustering without considering molecular symmetries is then performed on these rotated projection directions. In contrast to the clustering method we propose, the true topology induced by the molecular symmetry is neglected. The closer the projection direction is to the edge of the fundamental domain, the more severe the consequences (i.e., error) of neglecting the molecular symmetry will be.

## 7 Discussion about potential applications

In the present work, we have enabled the statistics and clustering method for both spatial rotations and projection directions considering molecular symmetry, by developing an approximation algorithm for calculating the mean and the variance. The proposed statistics and clustering method have the potential to be adopted in a wide range of applications.

A quaternion-assisted angular reconstruction algorithm[32, 33] contributes to solving a series of structures at 10-30Å resolution in 90s[34]. The key step in this algorithm is solving the absolute orientation problem[30, 31], which is related to the determination of the mean of a series of spatial rotations. With this present work, it becomes feasible to solve such problems in the context of considering molecular symmetry. This might lead to rethinking the possibility of adopting a quaternion-assisted angular reconstruction algorithm in state-of-the-art cryoEM reconstruction. Furthermore, the statistics of spatial rotations may contribute to the adoption of advanced statistical method, such as particle filter[77, 78, 8] or Kalman filter algorithm[79, 80, 81], in cryoEM image processing.

For the statistics of spatial rotations, they may be adopted to resolve continuous structural changes by manifold embedding[82, 83]. This is because in such methods, images are grouped by projection directions for the purpose of mapping the continuum of the states of a molecule in each projection direction. Moreover, they can be used for analyzing molecule heterogeneity and misalignment. As an example, 124,810 single particle images of a Tc complex are classified by projection directions when *C*_5_ symmetry is considered (Figure 2). The orientation heterogeneity of the tail exists, and approximately 12.5% images are misaligned. However, a 3.7Å resolution structure can be obtained from this dataset.

**Figure 2:**
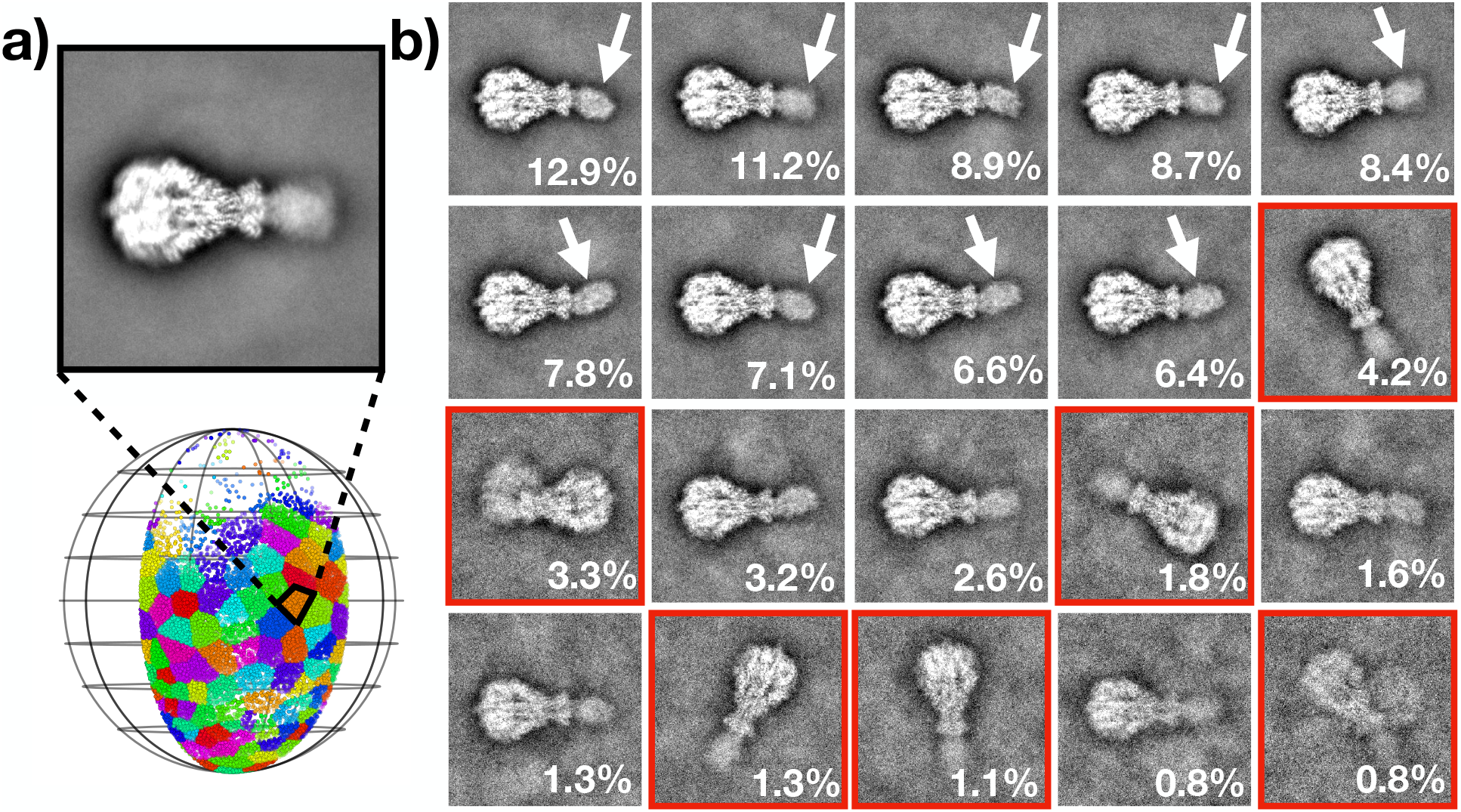
Projection direction clustering of a Tc complex. **a**, 124,810 single particle images of a Tc complex are classified by their projection directions into 100 classes, when *C*_5_ molecular symmetry is taken into consideration. Classes are distinguished by colors. The images from a selected class (orange) are averaged and then displayed. **b**, The images from the selected class are further classified into 20 classes. The orientation heterogeneity of the tail is indicated by an arrow. The misaligned images are boxed by red lines.

In summary, the statistics tools developed in this work form a system for analyzing spatial rotations and projection directions in cryoEM when molecular symmetry is considered. They provide more freedom and efficiency when processing and understanding images in cryoEM. We have realized the proposed methods in the open-source python package pySymStat, and it can be easily adopted or integrated in other algorithms or software.

## Acknowledgements

This work was supported by fund from Advanced Innovation Center for Structural Biology (to M.H.), Yau Mathematical Sciences Center (to H.L.), and grant TH-533310008 of Tsinghua University (to H.L.). The authors are grateful to Dr. Xueming Li for providing the Tc complex dataset, and to Dr. Yichen Zhou for revising the manuscript.

## Contributions

M.H., Q.Z. and H.L. initialized the project and designed the methods. Q.Z. and H.L. clarified mathematical issues and derived formulas. M.H. and Q.Z. wrote pySymStat and validated its performance. M.H. wrote the manuscript.

## Appendix I The sufficient condition of two spatial rotations being equivalent considering molecular symmetry

In this section, we prove the following statement. For a molecule with *G* symmetry, the spatial rotation **q** is equivalent to **q** ⊗ **g**_*m*_ for any **g**_*m*_ in the molecular symmetry group *G*. This statement can be paraphrased as follows. For two spatial rotations **q**_1_ and **q**_2_, the sufficient condition of **q**_1_ and **q**_2_ being equivalent considering molecular symmetry *G* is that there exists an element **g**_*m*_ in *G* such that **q**_1_ = **q**_2_ ⊗ **g**_*m*_.

A coordinate in the map is denoted by 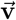. The value at coordinate 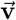 in this map is denoted by 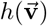. As this map has molecular symmetry *G*,

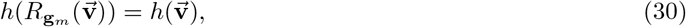

for all **g**_*m*_ in *G*.

The value at the coordinate 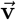 in the rotated map with a spatial rotation **q** is denoted by *h*_**q**_. By the definition of the spatial rotation,

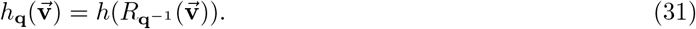

The sufficiency is proved as follows. As there exists an element **g**_*m*_ in *G* such that **q**_1_ = **q**_2_ ⊗ **g**_*m*_, it can be concluded that

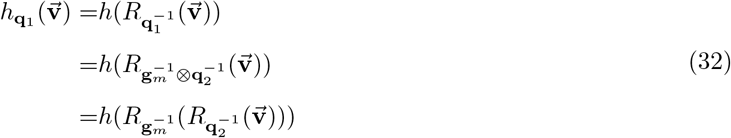

By Equation (30), it can be further concluded that

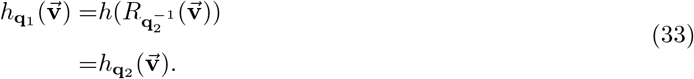

As stated in Equation (33), the **q**_1_ rotated map is identical to the **q**_2_ rotated map. Hence, **q**_1_ is equivalent to **q**_2_. Therefore, we have proven the sufficiency of the condition.

## Appendix II The sufficient condition of two projection directions being equivalent considering molecular symmetry

In this section, we continue to use the notations and conclusions presented in Appendix I. Next, we prove the following statement. For a molecule with *G* symmetry, the projection direction 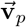 is equivalent to 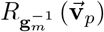 for any **g**_*m*_ in a molecular symmetry group *G*.

By the concept of the projection image set defined by Equation (4), the equivalence between two projection directions can be defined by the equivalence between the two sets of projected images. As the projection operator 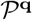 is defined by the operation of projecting along *Z* axis after after rotating the map *V* by spatial rotation **q** and the map *V* has molecular symmetry *G*, it stands that

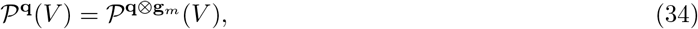

for any spatial rotation **q** and any element **g**_*m*_ ∈ *G*. Therefore, by definition of the projection image set, i.e., Equation (4), it can be concluded that

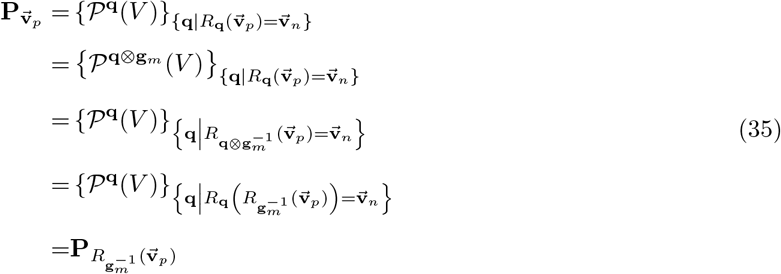

Therefore, we have proven the sufficiency of the condition.

## Appendix III Proof of well-definedness of the distance between two spatial rotations considering molecular symmetry

In this section, Equation (7) is proved to be a well-defined distance definition. The three requirements of distance definition in mathematics, i.e., the symmetry property, the positive definiteness property, and the triangle inequality property, are proven to be satisfied for Equation (7).

The symmetry property is such that

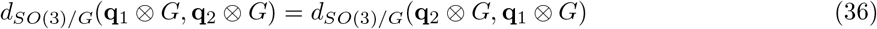

holds for all 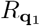, 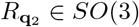. It is proven as follows.

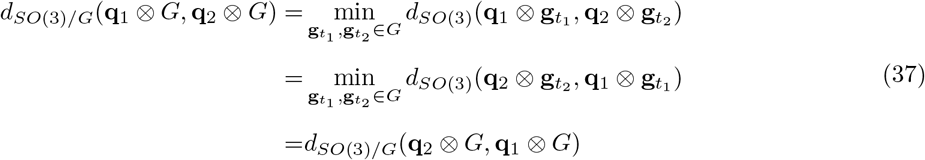

as 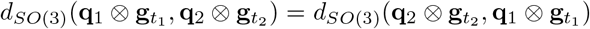 holds, because *d*_*SO*(3)_ is also a distance, therefore it has the symmetry property.

The positive definiteness property contains two parts. The first part is that

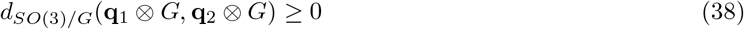

holds for all 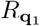, 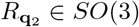, which can be derived by *d*_*SO*(3)_(**q**) ≥ 0 for all *R*_**q**_ ∈ *SO*(3). The second part is that

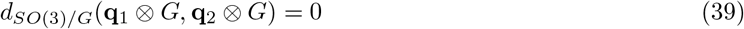

if and only if **q**_1_ ⊗ *G* = **q**_2_ ⊗ *G*. This statement is proven as follows. When **q**_1_ ⊗ *G* = **q**_2_ ⊗ *G*, there exist 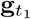 and 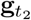 such that 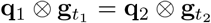. Thus, 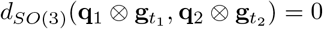, as *d*_*SO*(3)_ also has the positive definiteness property. Therefore, *d*_*SO*(3)*/G*_(**q**_1_ ⊗ *G,* **q**_2_ ⊗ *G*) = 0. Conversely, when *d*_*SO*(3)*/G*_(**q**_1_ ⊗ *G,* **q**_2_ ⊗ *G*) = 0, there exist 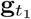 and 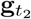 such that 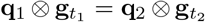. If **q**_1_ ⊗ *G* ≠ **q**_2_ ⊗ *G*, without losing generality, there exists 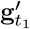 such that 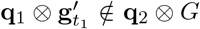. Therefore, 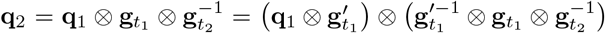 implies that 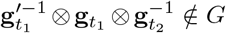, which conflicts with the property of the group *G*. Thus, **q**_1_ ⊗ *G* = **q**_2_ ⊗ *G*.

The triangle inequality property is that

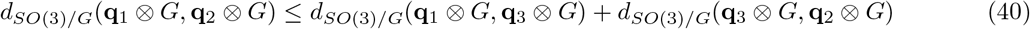

holds for all 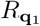, 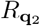, 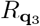 in *SO*(3). It is proven as follows.

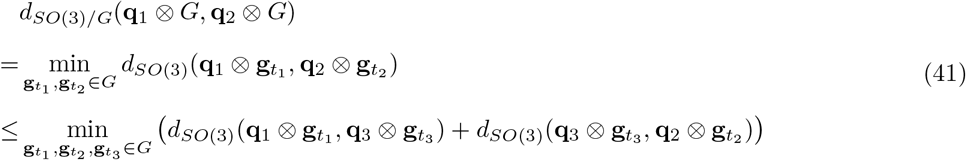

as 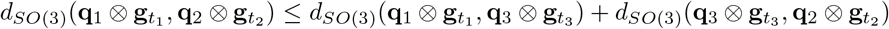 holds for any 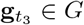, because *d*_*SO*(3)_ also has the triangle inequality property.

Moreover, distance in *SO*(3), i.e., *d*_*SO*(3)_, is invariant under left and right multiplication[1]. In other words, *d_SO_*_(3)_ has the following properties,

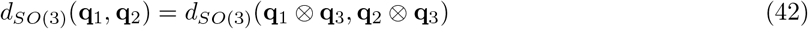

and

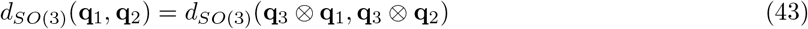

for any 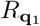, 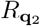 and 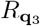 in *SO*(3). Therefore, Equation (41) can be further converted into

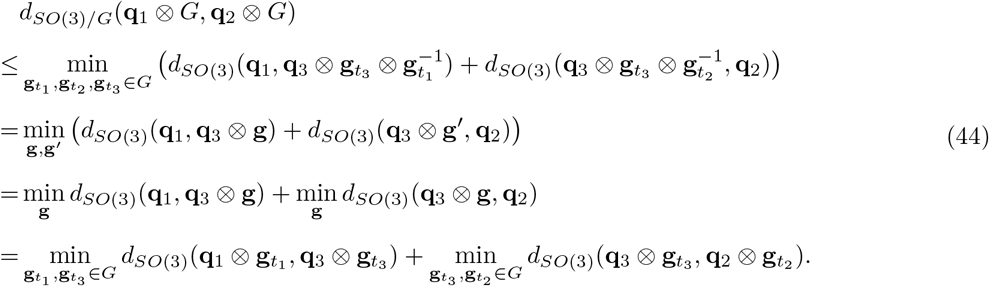

Therefore, we have proven the suitability of Equation (7) as a distance definition.

## Appendix IV Proof of well-definedness of distance definition between two projection directions considering molecular symmetry

In this section, Equation (8) is proven to be a well-defined distance definition. Three requirements of distance definition in mathematics, i.e., the symmetry property, the positive definiteness property and the triangle inequality property, are proven to satisfy Equation (8).

As the distance definition between two projection directions considering molecular symmetry, i.e., Equation (8), follows the same structure of the definition between two spatial rotations considering molecular symmetry, i.e., Equation (7), the symmetry property, the first part of the positive definiteness, and the triangle inequality property can be proven with the same procedure in Appendix III.

The second part of the positive definiteness is that

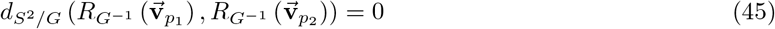

if and only if 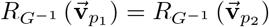. This statement is proven as follows. When 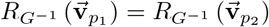, there exist 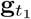 and 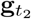 such that 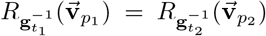. As 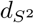 also has positive definiteness property 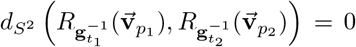. Therefore, 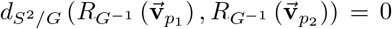 Conversely, when 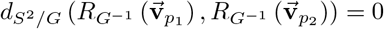 there exist 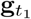 and 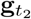 such that 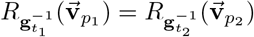. If 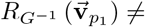 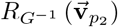, without losing generality, there exists 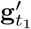 such that 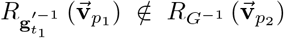. Therefore, 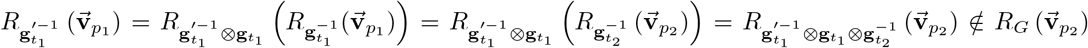, which conflicts with the property of the group *G*. Thus, 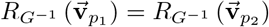.

Therefore, we have proven the suitability of Equation (8) as a distance definition.

## Appendix V Approximation of the mean of spatial rotations by representatives selected in variance determination

In this section, we state that it is rational to approximate the mean of spatial rotations, i.e, Equation (11), by Equation (13).

By observing the definition of mean and the definition of variance of spatial rotations, i.e., Equation (11) and (9), respectively, one can find that the spirit of the selection of 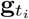 for the mean and the variance are the same. In other words, 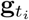 is selected for each **q**_*i*_ to make 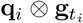 as close as possible overall. Therefore, it is natural to replace the 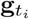 in the definition of the mean by the 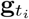 in the definition of the variance. Thus, Equation (13) is a rational approximation of Equation (11).

## Appendix VI Approximation of mean of projection directions by representatives selected in variance determination

In this section, we state that it is rational to approximate the mean of projection directions, i.e., Equation (12), by Equation (15).

As the relation between the mean and the variance of spatial rotations is identical to those of the projection directions, the rationality of approximating Equation (12) by Equation (15) can be explained by using the same procedure as in Appendix V.

## Appendix VII Derivation of representative determination of spatial rotations into a NUG form

In this section, we prove that Equation (14) can be derived into a NUG form, i.e., Equation (18).

Distance in *SO*(3), i.e., *d*_*SO(3)*_, is invariant under left and right multiplications[1], i.e., Equation (42) and Equation (43). Thus, Equation (14) can be derived as

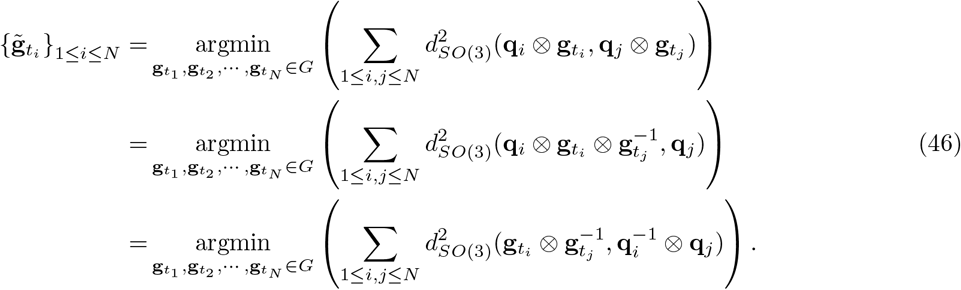

By assigning 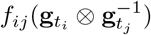 as Equation (19), we have proven that Equation (14) can be derived into a NUG form, i.e., Equation (18).

## Appendix VIII Derivation of representative determination of projection directions into a NUG form

In this section, we prove that Equation (16) can be derived into a NUG form, i.e., Equation (18).

The distance in *S*, i.e., 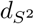 is invariant under spatial rotation. In other words, 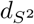 has the following property,

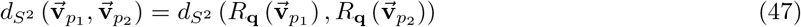

for any projection directions 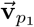 and 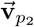, and any spatial rotation **q**.

Thus, Equation (16) can be derived into

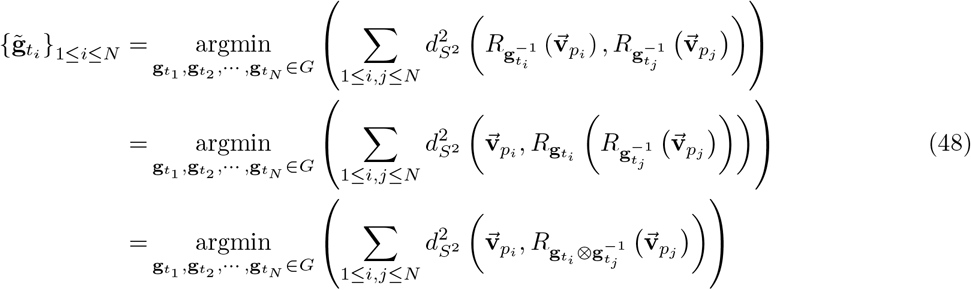

By assigning 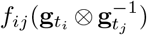 as Equation (20), we have proven that Equation (16) can be derived into a NUG form, i.e., Equation (18).

## Appendix IX Fourier expansion of a NUG, and a matrix form

In this section, we prove that Equation (18) is equivalent to Equation (23).

As *G* is a molecular symmetry group, *G* is compact. By the Peter-Weyl Theorem[2], the generalized Fourier expansion of 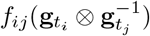 is

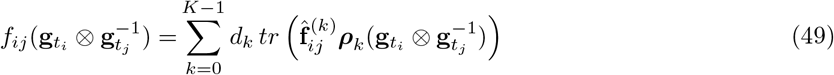

where 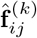 is determined by Equation (21).

Thus, Equation (18) can be derived into

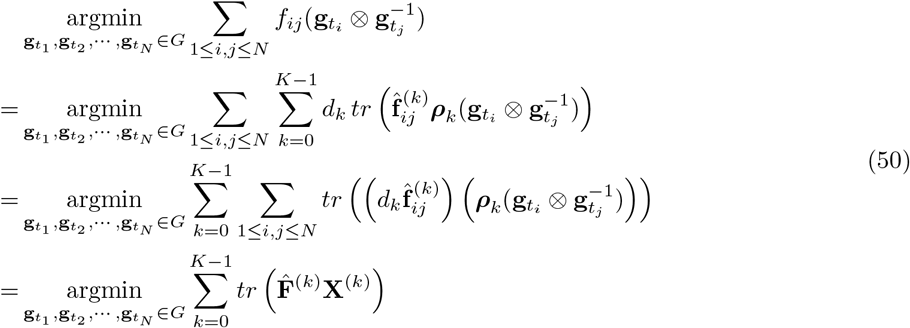

where 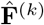 is determined by Equation (24) and **X**^(*k*)^ is determined by Equation (26). Hence, we have proven that Equation (18) is equivalent to Equation (23).

## Appendix X SDP relaxation of the matrix form of the NUG problem

The variable matrices {**X**^(*k*)^}_1≤*k*≤*K*_ is defined by Equation (26), in the matrix form of the NUG problem. However, the constraints of such matrices are in general non-convex. For the purpose of convex optimization, the constraints of {**X**^(*k*)^}_1≤*k*≤*K*_ are relaxed by the method proposed in[3]

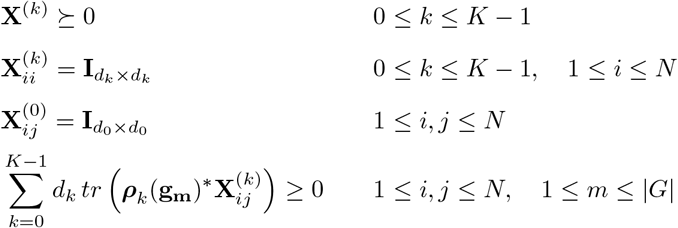

where **X**^(*k*)^ ≥ 0 stands for that {**X**^(*k*)^}_1≤*k*≤*K*_ are semi-definite positive (SDP) matrices.

Therefore, the NUG problem is relaxed into a SDP problem, as

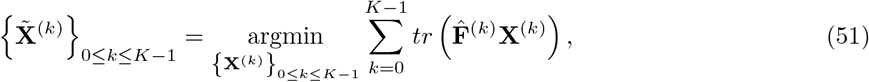

where {**X**^(*k*)^}_1≤*k*≤*K*_ are subject to the above relaxed constraint. The solution matrices 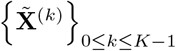 are in the convex hull of the matrices defined in Equation (26).

## Appendix XI Partial cost

Solving 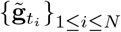 minimizes the cost function Equation (18). Thus, during the sequential recovery of representatives from the solution of the SDP problem, the partial cost must be calculated at each recovery step for the purpose of determining the keeping trials. As in Equation (18), the cost 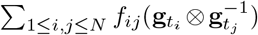 is the summation of *f_ij_* of all relations 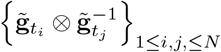. Therefore, the partial cost can be calculated as

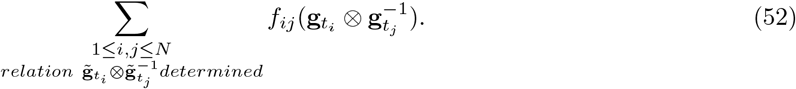

In other words, the partial cost is the summation of *f*_*ij*_ of all determined relations.

## Appendix XII Design of an intuitive approach

In the section, an intuitive approach for solving Equation (14) is described.

The principle of obtaining 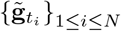 from Equation (14) is that a representative 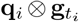 is chosen from each set of equivalent spatial rotations **q**_*i*_ ⊗ *G*, such that all representatives are as close as possible overall. In this intuitive approach, this principle is realized as follows.

First, a spatial rotation 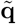 is selected as follows. For each **q**_*i*_ ⊗ *G* from {**q**_*i*_ ⊗ *G*}_1≤*i*≤*N*_, the overall distance between it and the other set of equivalent spatial rotations is calculated as

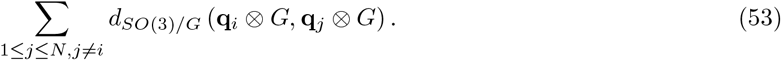

The set of equivalent spatial rotations with minimum such distance can be considered at the “center” of {**q**_*i*_ ⊗ *G*}_1≤*i*≤*N*_. Therefore, the spatial rotation 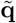 is selected as an element of the “center” set of equivalent spatial rotations.

Then, from each set of equivalent spatial rotations **q**_*i*_ ⊗ *G*, the one closest to the “center” 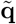, i.e., 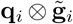, is selected as the representative. Therefore, 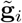 is determined for a spatial rotation set **q**_*i*_ ⊗ *G*.

By this means, a representative 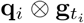 is chosen from each set of equivalent spatial rotations **q**_*i*_ ⊗ *G*. Such a representative is closest to the “center” 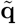. The principle of obtaining 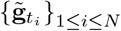 from Equation (14) is realized to some extent.

